# Incidental Temporal Binding in Rats: A Novel Behavioral Task Relevant to Episodic Memory

**DOI:** 10.1101/2022.08.30.505895

**Authors:** Dominika Radostova, Daniela Kuncicka, Branislav Krajcovic, Lukas Hejtmanek, Tomas Petrasek, Jan Svoboda, Ales Stuchlik, Hana Brozka

**Affiliations:** Insitute of Physiology, Czech Academy of Sciences, Videnska 1083, 142 20 Prague, Czechia; Second Faculty of Medicine, Charles University, V Uvalu 84, 150 06 Prague, Czechia; Third Faculty of Medicine, Charles University, Ruska 2411, 100 00 Prague, Czechia; National Institute of Mental Health, Topolova 748, 250 67 Klecany, Czechia

**Keywords:** Associative learning, episodic-like memory, temporal binding, active avoidance, conditioning

## Abstract

We designed a behavioral task called One-Trial Trace Escape Reaction (OTTER), in which rats incidentally associate two temporally discontinuous stimuli: a neutral acoustic cue (CS) with an aversive stimulus (US) which occurs two seconds later (CS-2s-US sequence). Rats are first habituated to two similar environmental contexts (A and B), each consisting of an interconnected dark and light chamber. Next, rats experience the CS-2s-US sequence in the dark chamber of one of the contexts (either A or B); the US is terminated immediately after a rat escapes into the light chamber. The CS-2s-US sequence is presented only once to ensure the incidental acquisition of the association. The recall is tested 24 h later when rats are presented with only the CS in the alternate context (B or A), and their behavioral response is observed. Our results show that 59 % of the rats responded to the CS by escaping to the light chamber, although they experienced only one CS-2s-US pairing. The OTTER task offers a flexible high throughput tool to study memory acquired incidentally after a single experience. Incidental acquisition of association between temporally discontinuous events is highly relevant to episodic memory formation.

## 1. INTRODUCTION

Episodic memory is the ability to retain and recall knowledge of personally experienced past events [1]. These events are often composed of successive sub-events separated by time gaps and might be experienced as a single memory [2]. The ability to form associations between events, known as temporal binding, is likely an essential prerequisite for creating complex episodic memories [3]. Another aspect of episodic memories is that they are acquired incidentally, i.e., we remember events we did not intend to memorize. In an experimental setup, incidental memory can be tested in situations when subjects are unaware that they will be tested on recall [4–6]. We believe that the acquisition of episodic-like memory should not require conditioning or pre-training; moreover, evidence suggests that mechanisms of incidental memory acquisition might differ from acquisition with intent [7–10].

To understand the neural mechanism of episodic memory, it is vital first to elucidate the neural mechanisms of acquisition and retrieval of temporally bound sub-events. The first step towards this goal is to develop a valid and reliable behavioral task. Our goal was to develop a simple temporal binding task with a clear behavioral response and a balanced success ratio. We focused on a one-trial design to ensure that memory was acquired incidentally. To achieve this, we took advantage of two natural behavioral tendencies of rodents: rodents avoid brightly lit environments [11] and actively escape an immediate threat [12,13].

Our task, which we named One-Trial Trace Escape Reaction (OTTER), consists of three phases: habituation, pairing, and recall (Figure 1A–C). The purpose of habituation is to familiarize rats with a novel environment and to reduce their exploratory activity. Each rat is habituated to two similarly constructed environmental contexts: an oval-shaped context A and a slightly larger rectangular-shaped context B (Figure 1A). Both contexts consist of one dark and one light chamber and are separated by a partition with a rectangular opening. This design allows us to exploit the natural tendency of rodents to avoid bright light. Even if rats are free to move between both chambers, they strongly prefer the dark chamber. To olfactorily distinguish both contexts, context A is cleaned with an alcohol-based wash, while context B is cleaned using a vinegar-based wash.

**Figure 1.**
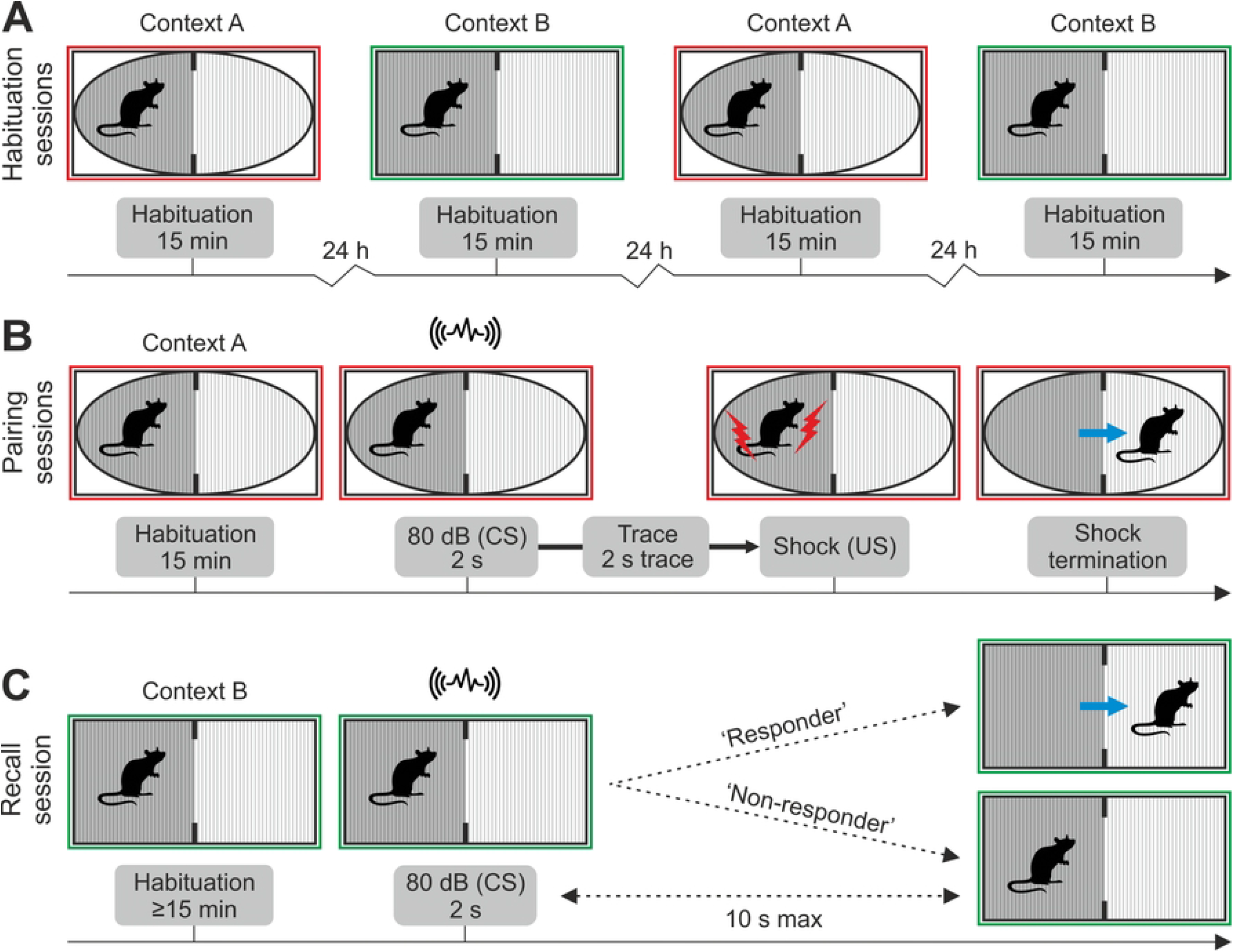
Schematic overview of One-Trial Trace Escape Reaction (OTTER). (**A**) A rat is initially habituated to environmental contexts A and B in alternating daily sessions. (**B**) During the pairing session, the rat hears a sound cue (CS) while in the dark chamber of one of the two contexts (context A or B); two seconds later, the rat receives a foot shock (US) that is terminated once the rat transfers to the light chamber. (**C**) The recall of CS-2s-US association is tested in the recall session, which occurs 24 hours later in the alternate context (context B or A). The CS is delivered when the rat settles in the dark chamber, and the rat’s reaction is observed. Upon hearing the CS, the rat either escapes into the light chamber (‘responder’) or stays in the dark chamber (‘non-responder’).

During the pairing session, rats experience two novel stimuli separated by a time gap (Figure 1B). Each rat is first allowed to explore the apparatus of context A or B exactly as it would during the habituation sessions. When the rat is settled in the dark chamber, it first hears a three-second acoustic cue (the conditioned stimulus CS), then receives an electric foot shock (unconditioned stimulus, US) two seconds after the CS stops. The US is terminated immediately after the rat escapes to the light chamber. This is the opportunity for the rat to incidentally associate the CS with the US (CS-2s-US) and learn that escape to the light chamber provides safety from the US.

The association between the CS and US is tested during the recall session 24 hours later (Figure 1C). Unlike in traditional active avoidance tasks, the recall in OTTER is tested in a different environmental context. Testing the recall in a different environmental context renders the association between the US and environmental context irrelevant; fear-related behavioral responses are therefore only attributable to the association between the CS and US. The recall session begins by placing the rat in the dark chamber of the environmental context other than the context of the pairing session (B or A). If at least 15 minutes elapsed and the rat rests in the dark chamber, the CS is delivered, and the rat’s reaction is observed. There are two possible reactions: either the rat escapes into the light chamber (‘responder’) or remains in the dark chamber (‘non-responder’). Here we present normative data from the OTTER task and discuss its strengths, limitations, and possible applications.

## 2. METHODS

### 2.1 Animals

Adult male Wistar rats (ENVIGO; 12–14 weeks old) were used in the experiment (n = 32). Upon arrival, the 28-day-old rats were housed in standard laboratory cages (50 × 25 × 25 cm), two animals per cage. Laboratory food and tap water were supplied *ad libitum*. The room where the animals were kept was ventilated with a constant temperature of 22 °C and 50% humidity. The rats were kept on a 12-hour light cycle, with light being turned on daily at 6 am.

Before the start of the OTTER task, all rats were handled by the experimenter for 3 minutes daily for four days. All experiments were conducted during the light phase of the day (9 am to 1 pm) because rats show lower locomotion during that time [14]. All animal procedures were approved by the Ethical Committee of the Czech Academy of Sciences and complied with the Animal Protection Act of the Czech Republic and EU directive 2010/63/EC.

### 2.2 Apparatus

Two modified TSE multi-conditioning shuttle boxes (TSE Systems GmbH, Germany) were used in the experiment. Each shuttle box consisted of two interconnected 24 × 47 cm chambers. The first chamber of both shuttle boxes was built from transparent acrylic glass (light chamber), while the second chamber was created using dark opaque acrylic glass (dark chamber). A dark opaque lid was used to cover the dark chamber, resulting in light intensity of less than 3 lx in the chamber’s center. The light chamber was left uncovered; moreover, we added another light source to reach a light intensity of 1090 lx in the chamber’s center. Intense light is highly uncomfortable for rodents [15], and aversively motivated rats spend most of their time in the dark chamber. The chambers were separated by a custom-made partition with a wide opening (4 × 40 cm central opening, custom-made) made of black acrylic glass.

The shuttle boxes were soundproofed and equipped with a speaker. Once triggered by the TSE software, the speaker delivered a 2400 Hz sound cue. The sound cue was delivered at 80 dB SPL intensity for 2 seconds. The walls of both chambers were equipped with infrared devices that registered the animal’s location within the apparatus. The floor of both chambers consisted of a metallic grid with 0.5 cm diameter metal rods spaced 1.5 cm apart. When prompted, the metal rods delivered a 1.0 mA pulsatile electric stimulus with a 400 ms period (a 200 ms, 1 mA stimulus followed by 200 ms no stimulus) to the animal in the dark chamber.

The two TSE multi-conditioning shuttle boxes were visually and olfactorily distinct so that one shuttle box served as environmental context A and the other as environmental context B. The walls of the light chamber were decorated with an aquarium scene on a circular insert in context A, while in context B, the walls of the light chamber were decorated with black stripes. Context A was cleaned with an alcohol-based wash, while context B was cleaned with a vinegar-based wash.

We presume the OTTER task can be conducted using any similar apparatus where the above-described general principles are adhered to and will deliver comparable results to those using TSE shuttle boxes.

### 2.3 Habituation, Pairing, and Recall

Each rat was individually habituated to the environmental context A and B twice in a series of four 15-minute habituation sessions; only one habituation session took place every day, and the sessions in contexts A and B alternated daily. At the start of every habituation, each rat was placed in the dark chamber and was left to explore both chambers freely.

Following habituation, rats were conditioned to CS-2s-US pairing. The pairing took place in the same context as the first habituation session: rats first exposed to context A experienced CS-2s-US pairing in context A and vice versa for context B. The beginning of the pairing session closely resembled the habituation session, as rats were allowed to move freely through the apparatus for 15 minutes. At this point, rats did not transfer between the chambers at all or transferred only seldom. Following the 15-minute interval, a to-be-conditioned stimulus ‒ a three-second sound cue ‒ was delivered. We advise that the CS should be delivered with caution, as delivering the CS at an inappropriate moment might hamper the CS-2s-US acquisition. The rat must be located in the dark chamber, resting and not facing the opening in the partition between chambers (to avoid bias toward escaping through the ‘door’). Two seconds after the CS stopped, an electric foot shock was delivered (US) to the rat by the metallic floor grid. The US was automatically terminated if the rat’s position was registered in the light chamber or if the rat did not leave the dark chamber in 20 seconds. Rats that did not escape to the light chamber were excluded and did not proceed to the recall session. Rats that escaped were returned to their home cage immediately after the escape and were left undisturbed for the next 24 hours, after which they were tested for recall.

The recall of the CS-2s-US pairing took place in the alternate context, i.e., if the pairing took place in context A, the recall was tested in context B. Recall session resembled the pairing session with the exception that the US was not delivered. The CS was delivered no sooner than after 15 minutes and only if the rat rested in the dark chamber. Following the CS delivery, the rat’s response was observed. Rats that escaped to the light chamber within 10 seconds of the CS start were considered ‘responders,’ while those that remained in the dark chamber were considered ‘non-responders.’

### 2.4 Statistics

IBM SPSS Statistics (version 25.0, IBM Corp., 2017) was used to analyze behavioral data. More than half of the data did not meet parametric assumptions; therefore, we used non-parametric tests in all statistical analyses (Mann-Whitney test, Friedman test, Wilcoxon-signed rank test, Kruskal-Wallis test, Fisher’s exact test). The significance threshold was set to p = 0.05.

### 2.5 Data visualization

Data visualizations were created in IBM SPSS Statistics (version 25.0, IBM Corp., 2017), Corel, and R using the visualization library ggplot2 [16]. Heatmaps were obtained using the two-dimensional kernel density estimation function, kde2d, from the MASS library [17].

## 3. RESULTS

### 3.1 Habituations

During the four 15-minute habituation sessions in environmental contexts A and B, rats preferred the dark chamber in both contexts (Figure 2A). We found that rats transferred significantly less to the light chamber in context B than in context A during the first habituation session (Mann-Whitney test: hab 1: U = 55.00, *p* = 0.008). The difference between number of transfers to the light chamber in context A and B was not significant during any other habituation session (Figure 2B) (Mann-Whitney test: hab 2: U = 112.50, *p* = 0.775; hab 3: U = 72.00, *p* = 0.057; hab 4: U = 116.50, *p* = 0.899). For plots of movement through the apparatus by the individual rats during the 15-minute habituation sessions, see Supplementary figure 1. We found no significant difference in the time spent in the dark chamber (Supplementary figure 2) (Mann-Whitney test: hab 1: U = 94.00, *p* = 0.315; hab 2: U = 96.50, *p* = 0.361; hab 3: U = 97.00, *p* = 0.373; hab 4: U = 88.00, *p* = 0.213) in context A and B during any of the four habituation sessions.

**Figure 2.**
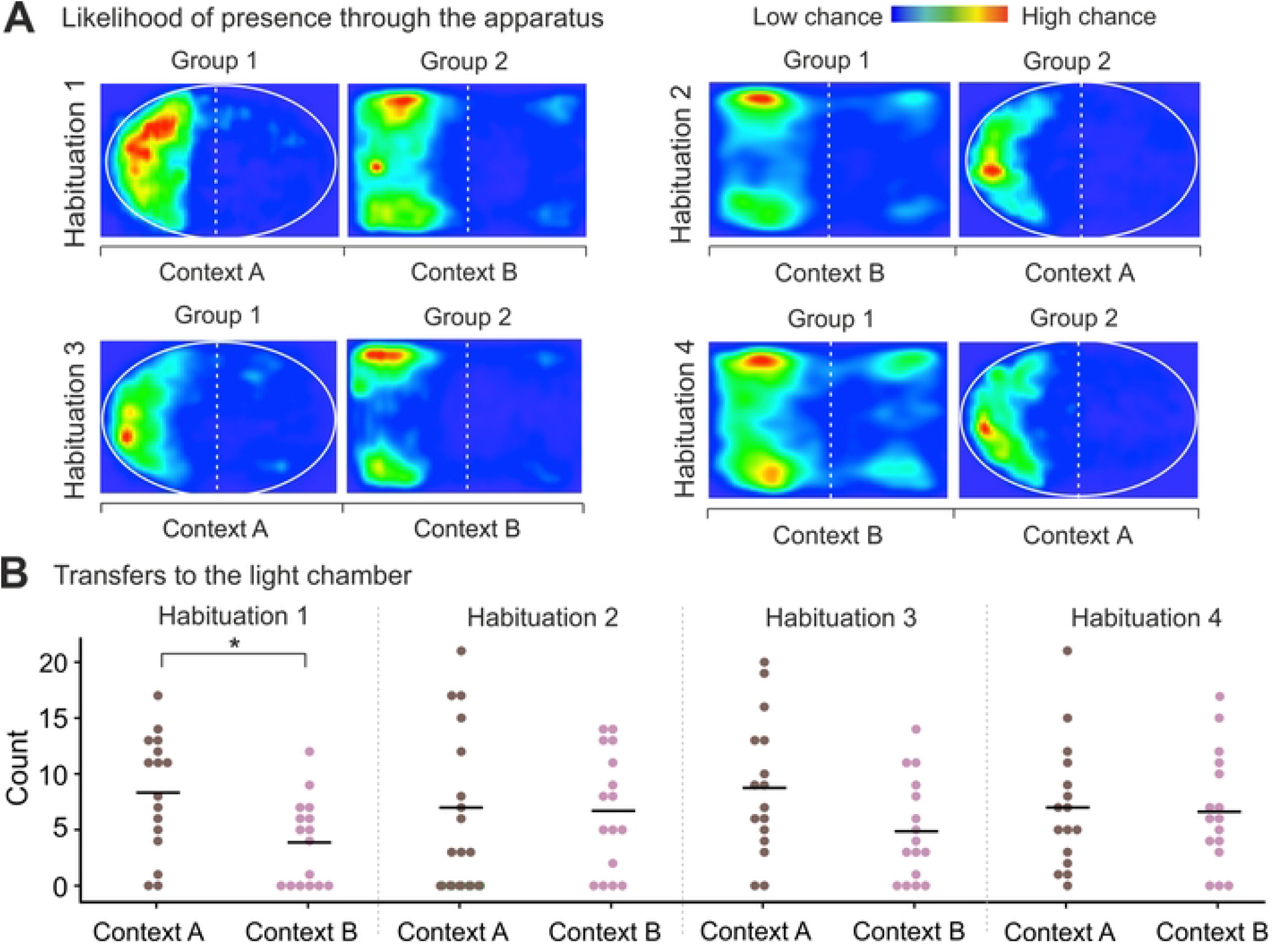
Chamber preference and transfers to the light chamber during 15 min habituation sessions. (**A**) Where animals were likely to be during each habituation session; blue indicates minimal presence, and red signifies a frequent stay. Rats were most often present in the dark chamber of contexts A and B during each session; the time spent in the dark chamber did not significantly differ between contexts A and B during habituation sessions. (**B**) Transfers to the light chamber in contexts A and B during habituation sessions. Rats transferred to the light chamber significantly less in context B than in context A during the first habituation session (p < 0.01), but not during any other habituation session.

**Supplementary figure 1. Movement of each rat through the apparatus during four 15-minute habituation sessions.**Each horizontal line corresponds to one animal. Black represents a stay in the dark chamber; gray represents a stay in the light chamber. Symbols “A” and “B” on the vertical axis correspond to the environmental context of the rat’s habituation. Context A was oval-shaped and cleaned with an alcohol-based wash, while context B was rectangular-shaped and cleaned with a vinegar-based wash.

**Supplementary figure 2. Time spent in chambers of contexts A and B by rats during habituation sessions.** Each rat (N = 32) received two habituation sessions in each context in an alternating manner. Starting context was chosen randomly for each rat. Rats preferred the dark chamber and spent very little time in the light chamber in both contexts across habituation sessions. The difference in the time spent in the dark chamber was significantly higher than the time spent in the light chamber in each context and during all five habituation sessions (Mann-Whitney test: Hab 1 context A: U = 0.000, *p* = 0.000; Hab 1 context B: U = 0.000, *p* = 0.000; Hab 2 context A: U = 1.000, *p* = 0.000, Hab 2 context B: U = 0.000, *p* = 0.000; Hab 3 context A: U = 1.000, *p* = 0.000, Hab 3 context B: U = 6.000, *p* = 0.000; Hab 4 context A: U = 1.000, *p* = 0.000, Hab 4 context B: U = 0.000, *p* = 0.000. Error bars indicate SEM, * indicates *p* <0.05.

As each rat was habituated to the same context twice, we also assessed if the number of transfers to the light chamber and the time spent in the dark chamber differed between these two habituation sessions. We found that in group 1 (first habituation in context A) the number of transfers to the light chamber did not significantly change between the two sessions in neither context A (Wilcoxon signed-rank test: Z = -0.473, *p* = 0.658) nor in context B (Z = -0.199, *p* = 0.873). Similarly, we found no significant difference in the time spent in the dark chamber between the two sessions in neither context A (Wilcoxon signed-rank test: Z = -0.879, *p* = 0.379) nor in context B (Z = -0.314, *p* = 0.753) in group 1. In group 2 (first habituation in context B), we observed similar behavior: the number of transfers to the light chamber did not significantly change between the two habituations sessions in context A (Wilcoxon signed-rank test: Z = -0.063, *p* = 0.964) nor in context B (Z = -1.129, *p* = 0.283). Time spent in the dark chamber also did not significantly differ between the two sessions in the same context (Wilcoxon signed-rank test: Z = -0.549, *p* = 0.583; Z = -0.879, *p* = 0.379 for context A and context B, respectively).

### 3.2 Pairing sessions

At the beginning of the pairing session, 24 rats were randomly selected to be presented with CS-2s-US (test group) and 8 rats to be presented with CS only (control group). One rat in the test group was excluded at the beginning of the pairing session as it lingered in the light chamber, and CS-2s-US could not be presented. All other test group rats (N = 23) were presented with CS-2s-US and escaped to the light compartment within 20 seconds. The average latency to escape was 10.0 ± 0.9 seconds (SEM) from the CS onset. Rats that responded to CS during the recall the next day escaped the US on average slightly faster than future non-responders (9.3 s compared to 11.0 s) (Figure 4C); however, this difference was not statistically significant (Mann-Whitney test: U = 29.000, *p* = 0.601). None of the control rats (N = 8) transferred to the light chamber upon hearing the CS.

During the habituation session preceding recall, rats transferred to the light chamber significantly less than during habituation before CS-2s-US pairing (Wilcoxon signed-rank test: Z = -2.762, *p* = 0.004) (pairing session habituation Mdn = 5.5, recall session habituation Mdn = 3) (Figure 3A). Although rats appeared to spend more time in the dark chamber during habituation preceding recall (Mdn = 843.8 s) than during habituation before pairing (Mdn = 814.4 s), the difference was not significant (Wilcoxon signed-rank test: Z = -0.843, *p* = 0.410) (Figure 3B).

**Figure 3.**
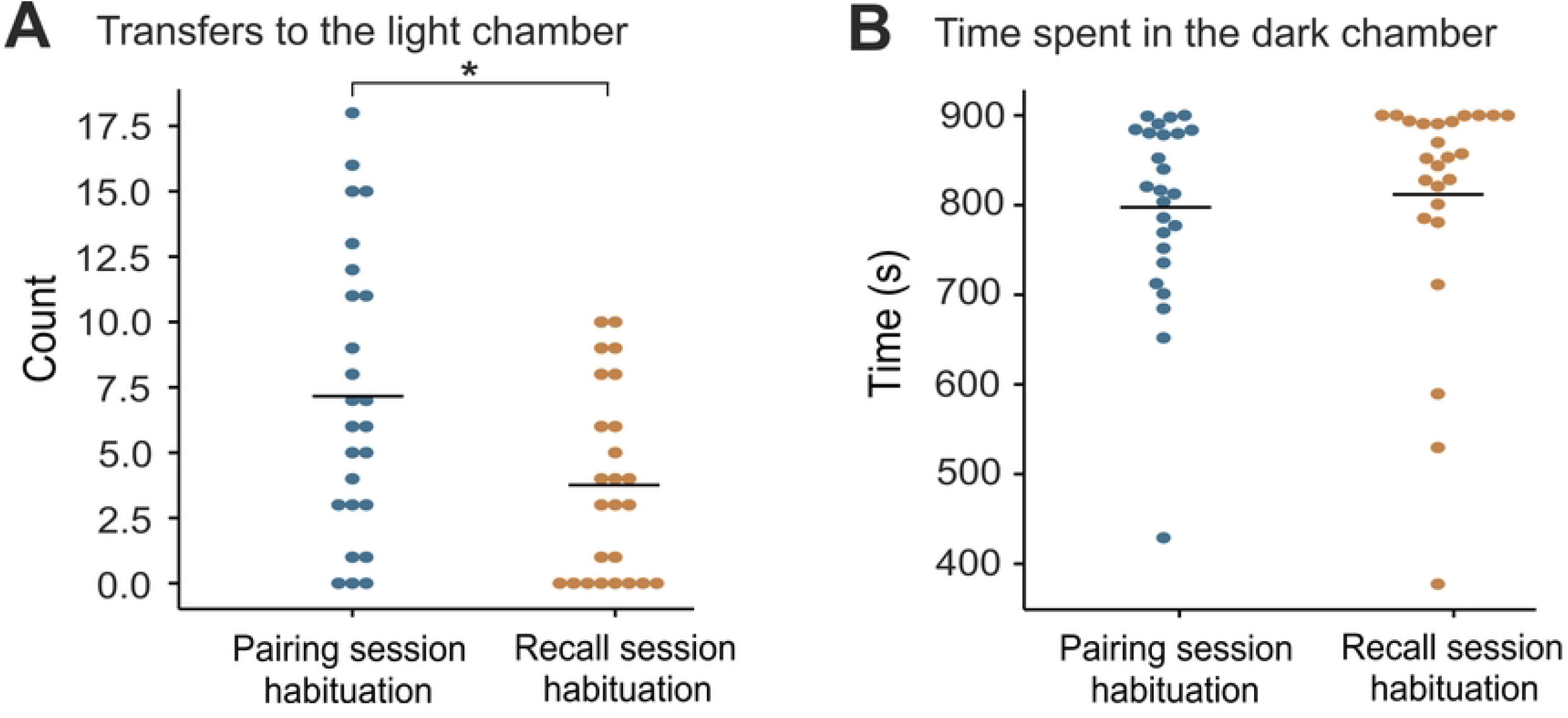
Comparison of transfers to the light chamber and time spent in the dark chamber during pairing and recall session. (**A**) Transfers to the light chamber during pairing and recall session habituations (data from contexts A and B combined). Rats transferred to the light chamber significantly less during recall session habituation (p < 0.01). (**B**) Time spent in the dark chamber during pairing and recall session habituations (data from contexts combined). The difference in time spent in the dark chamber between these two sessions did not significantly differ.

### 3.3 Recall sessions

During the recall session, the CS was successfully presented to 17 rats from the test group and 8 animals from the control group; 6 rats from the test group were excluded from the experiment as they lingered in the light chamber during the time of the intended CS presentation (Figure 4A). In the test group, 59 % (N = 10) of rats moved to the light chamber within 10 seconds of the start of CS (Figure 4B). The average time to escape was 5.1 ± 0.9 seconds (SEM) from the CS onset (Figure 4D). None of the control rats (N = 8) transferred to the light chamber in response to the CS. The number of responders in the test group was significantly higher than in the control group (Fisher’s exact test: p (two-tailed) = 0.008). We found no statistically significant difference in the time spent in the dark chamber (Kruskal-Wallis test: H(2) = 1.674, p = 0.433) (Figure 4E) or in the number of transfers to the light chamber (Kruskal-Wallis test: H(2) = 1.384, p = 0.501) between responders, non-responders and controls during the habituation session before recall (Figure 4F).

**Figure 4.**
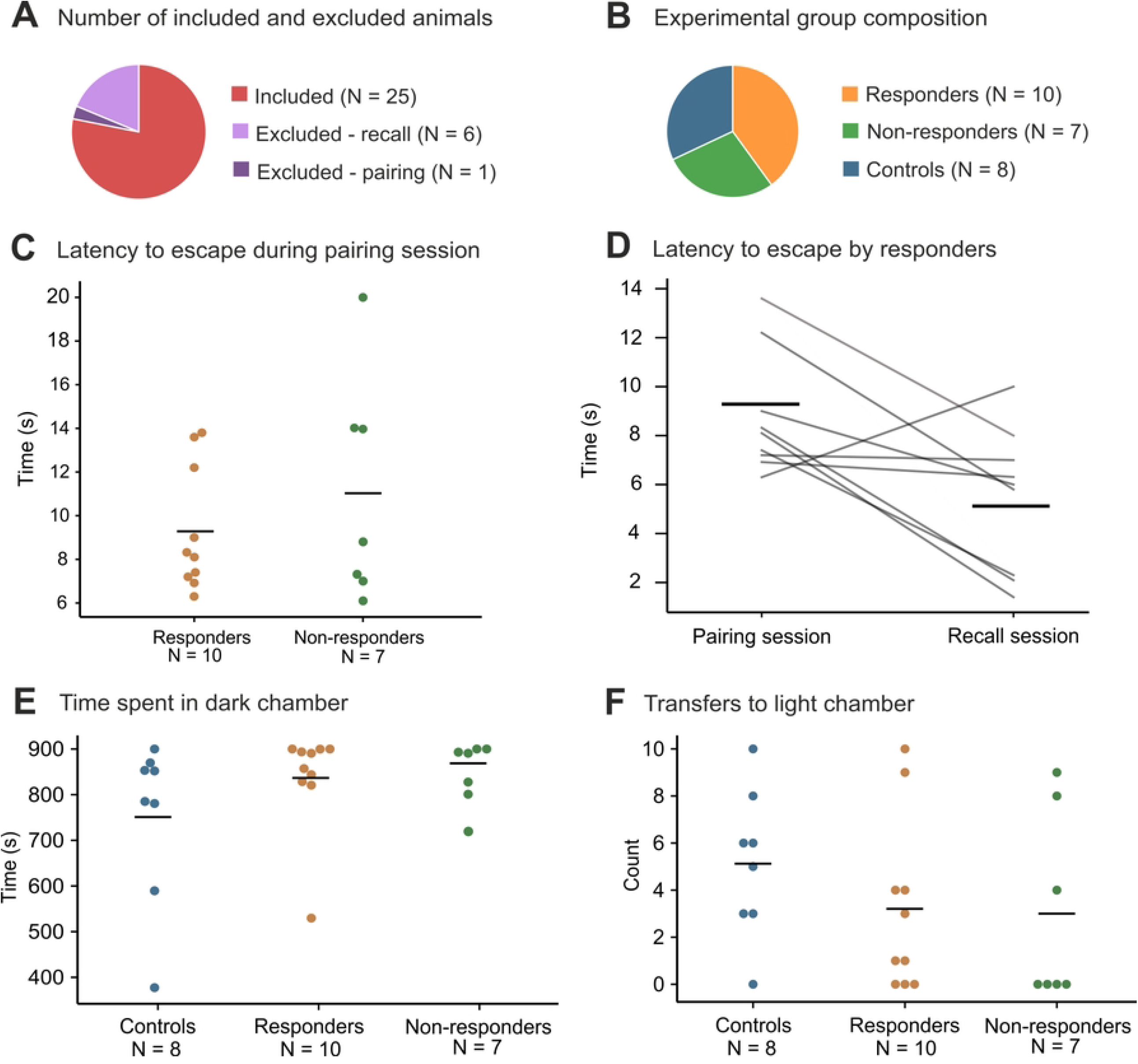
The OTTER task normative data. (**A**) The pie chart illustrates animals included in the study (N = 25) and animals excluded before the pairing (N = 1) and the recall session (N = 6). (**B**) Ratios of responders (N = 10), non-responders (N = 7) and controls (N = 8) in the group of 25 animals included in the study. Animals in the control group did not respond to CS during the recall phase. (**C**) Latencies to transfer to the light chamber during the pairing (since CS-2s-US start) by responders and non-responders. Non-responders transferred to the light chamber after 11.0 ± 1.9 seconds and responders after 9.3 ± 0.9 seconds. (**D**) Latencies to transfer to the light chamber by responders during pairing (since CS-2s-US start) and recall (since CS start; CS length was 2 seconds). Responders transferred to the light chamber after 9.3 ± 0.9 seconds during the pairing and after 5.1 ± 0.9 seconds during the recall. (**E**) Time spent in the dark chamber during the habituation preceding recall by non-responders, responders, and controls. (**F**) Transfers to the light chamber during the habituation preceding recall by non-responders, responders, and controls.

## DISCUSSION

We designed a behavioral task – OTTER – in which a single event is sufficient for an animal to incidentally acquire knowledge of a contingency existing over a time gap. The animal later actively exhibits this knowledge in a way that allows unambiguous evaluation of memory acquisition. The task utilizes species-specific preferences and naturally occurring behavior, which greatly reduces the amount of training needed and increases the task’s ecological validity. The OTTER task is a trace conditioning task that employs active avoidance to capture an important feature of naturalistic human episodic memory: the incidental memory acquisition of an event experienced only once.

The OTTER does not need pre-training or ‘priming’ the animals to anticipate a behaviorally relevant event. The pre-training in existing rodent paradigms can be twofold: shaping the desired response behavior or repeated exposure to stimuli. Both types of pre-training create an expectancy of contingency. For example, in trace fear conditioning, the CS-trace-US is presented more than once [18,19]. Although rodents may associate CS with US even after a single CS-trace-US exposure in trace fear conditioning, no observable variable could indicate the acquired knowledge. Freezing, an outcome measured in trace fear conditioning and similar tasks, is a response associated with an absence of an escape route [20] and might indicate behavioral despair as no action can avert the stressor. In this context, it is possible that freezing behavior emerges only after repeated CS-trace-US presentation. The OTTER task bypasses the potentially non-specific freezing response because a single pairing session results in a clear avoidance response during the recall session in rats.

In the OTTER task, active avoidance behavior offers an unambiguous binary measure of recall so that each animal can be confidently singled out as a ‘responder’ or ‘non-responder.’ In contrast, fear conditioning tests and novel object recognition tasks use a continuous variable as an indicator of learning, such as freezing or duration of exploration. When a continuous variable is used to classify animals into discrete groups, we first need to set a threshold value. Choosing the threshold value may prove challenging, and even then, it is often unclear how confident we can be about classifying animals just around the threshold values. Using a continuous outcome variable as a basis for classification might therefore provide unclear results. Rodent tasks with unambiguous output variables usually involve ‘declaring’ the knowledge by entering a correct place or pressing a correct button [5,21]. However, extensive pre-training is often required since the declarative behavior must be shaped first (usually approach behavior). The need for long pre-training to achieve declarative behavior precludes the study of incidentally acquired memories. In the OTTER task, the response is binary, and rats naturally exhibit the chosen declarative behavior without prior training.

To confidently ascribe the escape reaction to the CS-2s-US association, we must first establish an invariant baseline behavior — the animal has to stay in the dark chamber most of the time on its own accord. To this aim, we first explored the influence of rat strain on behavior within the same environmental context and found that Wistar rats display the most appropriate behavior for the OTTER task. In our experiments, Wistar rats spent most of the time in the dark chamber and transferred less frequently to the light chamber than Sprague-Dawley and Long-Evans rats (unpublished results). Second, we changed the size and shape of the opening in the partition between chambers which further reduced the time Wistar rats spent in the light chamber (unpublished results). By implementing these measures, rats seldom spontaneously move to the light chamber; hence it is very likely that when an animal ‘responds’ to the CS, it does so because it remembers the CS-2s-US association and not because of random exploration.

As long as the general principle of the OTTER task is adhered to, that is, controlling animal behavior by balancing conflicting species-specific behavioral tendencies, the OTTER task is flexible and can be embodied even by different physical instances. We are currently developing a second variant of the OTTER task with the working title ‘cold-OTTER.’ In the ‘cold-OTTER’ task, the invariant behavior is achieved by utilizing the rat’s and mice’s preference for warmth, which results in the avoidance of the cold sub-area of the apparatus. The flexible nature of the OTTER task allows adapting the task for different species and research contexts.

The OTTER task offers a high temporal precision of the recall event, making it an auspicious task for detailed studies of retrieval mechanisms. There is only a brief time window when an animal retrieves information and acts upon it. Such pinpointing of the recall event is difficult in tasks where a behavioral response is registered as a frequency of behavior during a time interval (freezing or exploration duration). The temporal precision of the recall event offered by the OTTER task can be especially advantageous if combined with high temporal resolution methods, such as electrophysiology [22] or calcium imaging [23]. The OTTER task might therefore serve as a valuable behavioral paradigm for a detailed study of neural mechanisms involved in episodic-like memory retrieval.

We consider OTTER highly relevant to episodic memory because the successful recall of CS-2s-US in OTTER meets several criteria of episodic memory: a) the memory was incidentally encoded [6], b) encoding occurred after a single exposure [24], c) there was no pre-training involved [25], d) rat behavior observed threshold retrieval dynamics [26], and e) rats were able to retrieve information flexibly in a different context [27]. Considering these five criteria, OTTER is a good model of several putative aspects of episodic memory. However, the OTTER task does not meet the episodic-like memory criterion of demonstration of what-where-when knowledge of past experiences [28] because the flight in response to the CS does not indicate if the rat remembers where and when it experienced CS-2s-US. Rather than puzzling over whether OTTER is ‘an episodic-like memory task’ or not, we find it more helpful to focus on the fact that the OTTER task captures several essential aspects of episodic memory and enables us to study them.

The OTTER task could be utilized to study the extinction of incidentally acquired memory based on a single exposure. This aspect is highly relevant to several neuropsychiatric disorders, especially post-traumatic stress disorder (PTSD). In this sense, the OTTER task could serve as an ecologically valid memory acquisition/extinction model of PTSD. We expect the extinction curve of CS-2s-US association could be influenced in both directions (faster/slower) by behavioral manipulations during or after the recall session.

As in any behavioral task, there are limitations to the OTTER task. First, the one-trial nature of the OTTER task precludes repeated measurements often required to accumulate sufficient amounts of data (e.g., in electrophysiology). This limitation stems from probing the incidental one-trial aspects of episodic-like memory and seems unavoidable. Second, it cannot be ruled out that ‘non-responders’ did form the CS-2s-US association but failed to act upon it. In assessing the recall, we rely on motoric output that is only indirectly related to the animal’s mental state. However, we found no significant difference in time spent freezing at the start of the recall session (Supplementary figure 3) and following the CS presentation (Supplementary figure 3) between ‘non-responders,’ ‘responders,’ and controls in our preliminary experiments. Our results suggest that ‘non-responders’ did not fail to learn that to avoid the US, they needed to escape to the light chamber but did not associate the US with either CS or the dark chamber at all.

**Supplementary figure 3. Freezing at the beginning of the recall session and after the CS presentation.** Freezing was assessed manually from video recordings by a blinded experimenter. As a freezing, we considered any lack of movement except for breathing. **(A)** Time spent freezing (s) during the first minute of the recall session. There was no significant difference in time spent freezing between ‘responders,’ ‘non-responders,’ and control rats (Kruskal-Wallis test: H = 6.436, p = 0.092). **(B)** Time spent freezing (s) during one minute after CS presentation. We found no significant difference in time spent freezing between ‘non-responders’ and control rats (Mann-Whitney test: U = 11.00, p = 0.831). This suggests that ‘non-responders’ did not recall the CS-2s-US association.

In conclusion, we designed a temporal binding task called OTTER that is adaptable, gives rapid results, and is easy to conduct. Due to the association of temporarily discontinuous events, the OTTER task can vastly promote our understanding of the neural mechanisms of temporal binding and possibly memory extinction. The behavioral response in the OTTER task is ecologically valid because it takes advantage of the natural behavioral tendencies of rodents. We demonstrated that rats could utilize the knowledge acquired from a single experience and use it to their advantage in a different context: rats demonstrated the same behavior that resulted in the termination of the unpleasant stimulus they experienced 24 hours earlier. We observed no ‘ceiling effect’ and a good balance between ‘responders’ and ‘non-responders’(close to 1:1), which may be valuable during tracing of neural changes in the early episodic-like memory acquisition. The OTTER task extends the current range of trace conditioning tasks, capturing the one-trial and incidental nature of encoding, and offers high temporal precision regarding when the memory recall occurred. Another notable advantage of the OTTER task is the binary outcome. The rat either crosses to the light chamber or does not; thus, there is no need to set an arbitrary threshold for the outcome variable. The OTTER task shares aspects with episodic memory due to its incidental, single-trial character with minimal training requirements.

## ACKNOWLEDGEMENTS

This work was supported by Czech Science Foundation (GACR) grant 20-00939S. Institutional support for IPHYS was provided by RVO: 67985823.

We would like to thank David Levcik for valuable feedback on the initial draft of the manuscript.

## AUTHOR CONTRIBUTIONS

Conceptualization, H.B., D.R., and B.K.; Methodology, D.R., and D.K.; Investigation, D.R. and D.K.; Resources, A.S, and J.S.; Writing – Original Draft, D.R., H.B, and B.K.; Writing – Review & Editing, H.B., D.K., B.K., T.P., and L.H.; Visualization, L.H., D.K., and H.B.; Supervision, H.B., and A.S.; Funding Acquisition, A.S.

## REFERENCES

1. Crystal JD. Animal models of episodic memory. Comp Cogn Behav Rev. 2018;13:105–22.

2. Tulving E. Episodic memory: From mind to brain. Annu Rev Psychol. 2002 Feb;53(1):1–25.

3. DuBrow S, Davachi L. Temporal binding within and across events. Neurobiol Learn Mem. 2016 Oct;134:107–14.

4. Zhou W, Crystal JD. Validation of a rodent model of episodic memory. Anim Cogn. 2011 May 17;14(3):325–40.

5. Zhou W, Hohmann AG, Crystal JD. Rats answer an unexpected question after incidental encoding. Current Biology. 2012 Jun 19;22(12):1149–53.

6. Zentall TR. Animals represent the past and the future. Evolutionary Psychology. 2013 Jul 1;11(3):573–90.

7. Rugg MD, Fletcher PC, Frith CD, Frackowiak RSJ, Dolan RJ. Brain regions supporting intentional and incidental memory: a PET study. Neuroreport. 1997 Mar 24;8(5):1283–7.

8. Trivedi MA, Murphy CM, Goetz C, Shah RC, Gabrieli JDE, Whitfield-Gabrieli S, et al. fMRI activation changes during successful episodic memory encoding and recognition in amnestic mild cognitive impairment relative to cognitively healthy older adults. Dement Geriatr Cogn Disord. 2008;26(2):123–37.

9. Wang WC, Giovanello KS. The role of medial temporal lobe regions in incidental and intentional retrieval of item and relational information in aging. Hippocampus. 2016 Jun;26(6):693–9.

10. Kuhnert MT, Bialonski S, Noennig N, Mai H, Hinrichs H, Helmstaedter C, et al. Incidental and intentional learning of verbal episodic material differentially modifies functional brain networks. PLoS One. 2013 Nov 18;8(11):e80273.

11. Keller FS. Light-aversion in the white rat. Psychol Rec. 1941 May 25;4(17):235–50.

12. Fanselow MS. Neural organization of the defensive behavior system responsible for fear. Psychon Bull Rev. 1994 Dec;1(4):429–38.

13. Wendt J, Löw A, Weymar M, Lotze M, Hamm AO. Active avoidance and attentive freezing in the face of approaching threat. Neuroimage. 2017 Sep;158:196–204.

14. Borbély AA, Neuhaus HU. Daily pattern of sleep, motor activity and feeding in the rat: Effects of regular and gradually extended photoperiods. J Comp Physiol. 1978;124(1):1–14.

15. Barker DJ, Sanabria F, Lasswell A, Thrailkill EA, Pawlak AP, Killeen PR. Brief light as a practical aversive stimulus for the albino rat. Behavioural Brain Research. 2010 Dec 25;214(2):402–8.

16. Wickham H. ggplot2: Elegant graphics for data analysis. Springer-Verlag New York; 2016.

17. Venables WN, Ripley BD. Modern Applied Statistics with S. Fourth edition. New York: Springer; 2002.

18. McEchron MD, Cheng AY, Gilmartin MR. Trace fear conditioning is reduced in the aging rat. Neurobiol Learn Mem. 2004 Sep;82(2):71–6.

19. Sharma V, Cohen N, Sood R, Ounallah-Saad H, Gal Ben-Ari S, Rosenblum K. Trace fear conditioning: procedure for assessing complex hippocampal function in mice. Bio Protoc. 2018 Aug 20;8(16):e2475.

20. Blanchard DC, Griebel G, Pobbe R, Blanchard RJ. Risk assessment as an evolved threat detection and analysis process. Neurosci Biobehav Rev. 2011 Mar;35(4):991–8.

21. Sato N. Episodic-like memory of rats as retrospective retrieval of incidentally encoded locations and involvement of the retrosplenial cortex. Sci Rep. 2021 Dec 26;11(1):2217.

22. Kim K, Vöröslakos M, Seymour JP, Wise KD, Buzsáki G, Yoon E. Artifact-free and high-temporal-resolution in vivo opto-electrophysiology with microLED optoelectrodes. Nat Commun. 2020 Dec 28;11(1):2063.

23. Scott BB, Thiberge SY, Guo C, Tervo DGR, Brody CD, Karpova AY, et al. Imaging cortical dynamics in GCaMP transgenic rats with a head-mounted widefield macroscope. Neuron. 2018 Dec;100(5):1045-1058.e5.

24. Morris RG. Episodic–like memory in animals: psychological criteria, neural mechanisms and the value of episodic–like tasks to investigate animal models of neurodegenerative disease. Philos Trans R Soc Lond B Biol Sci. 2001 Sep 29;356(1413):1453–65.

25. Binder S, Dere E, Zlomuzica A. A critical appraisal of the what-where-when episodic-like memory test in rodents: Achievements, caveats and future directions. Prog Neurobiol. 2015 Jul;130:71–85.

26. Eichenbaum H, Fortin NJ, Ergorul C, Wright SP, Agster KL. Episodic recollection in animals: “If it walks like a duck and quacks like a duck….” Learn Motiv. 2005 May;36(2):190–207.

27. Clayton NS, Bussey TJ, Dickinson A. Can animals recall the past and plan for the future? Nat Rev Neurosci. 2003 Aug;4(8):685–91.

28. Clayton NS, Bussey TJ, Emery NJ, Dickinson A. Prometheus to Proust: the case for behavioural criteria for ‘mental time travel.’ Trends Cogn Sci. 2003 Oct;7(10):436–7.

